# Genetic and shared couple environmental contributions to smoking and alcohol use in the UK population

**DOI:** 10.1101/555961

**Authors:** Toni-Kim Clarke, Mark J. Adams, David M. Howard, Charley Xia, Gail Davies, Caroline Hayward, Archie Campbell, Sandosh Padmanabhan, Blair H. Smith, Alison Murray, David Porteous, Ian J. Deary, Andrew M. McIntosh

## Abstract

Alcohol use and smoking are leading causes of death and disability worldwide. Both genetic and environmental factors have been shown to influence individual differences in the use of these substances. In the present study we tested whether genetic factors, modelled alongside common family environment, explained phenotypic variance in alcohol use and smoking behaviour in the Generation Scotland (GS) family sample of up to 19,377 individuals. SNP and pedigree-associated effects combined explained between 18% and 41% of the variance in substance use. Shared couple effects explained a significant amount of variance across all substance use traits, particularly alcohol intake, for which 38% of the phenotypic variance was explained. We tested whether the within-couple substance use correlations were due to assortative mating by testing the association between partner polygenic risk scores in 34,987 couple pairs from the UK Biobank (UKB). Only couples’ smoking status PRSs were significantly associated (b=0.01, S.E=0.005, p=0.02). However, an individual’s alcohol PRS was associated with their partner’s phenotype (b=0.04, S.E=0.007, p < 2 x 10^-7^). In support of this, G carriers of a functional *ADH1B* polymorphism (rs1229984), known to be associated with greater alcohol intake, were found to consume less alcohol if they had a partner who carried an A allele at this SNP. Together these results show that the shared couple environment contributes significantly to patterns of substance use. It is unclear whether this is due to shared environmental factors, assortative mating, or indirect genetic effects. Future studies would benefit from longitudinal data and larger sample sizes to assess this further.

## Introduction

Alcohol and tobacco have been used recreationally by humans across the world for centuries. The extent to which an individual uses alcohol and tobacco, and whether they use them at all, depends on individual genetics, environment, and cultural attitudes and the complex interactions between these factors. There has been extensive research into individual differences in alcohol and tobacco use, and the genetic component of these behaviours is well established. Heritability estimates range from 10 to 60% for alcohol use^1-3^, with alcohol use disorders tending to have higher estimates than for levels of consumption^3,4^. Similarly, smoking behaviours have a genetic component with the heritability estimates of nicotine dependence higher (60-70%) than tobacco use (ever vs never) (40-50%)^4-6^.

It is clear from heritability studies that a significant proportion of the phenotypic variance in individual differences comes from environmental, or other unmeasured sources, and measurement error. Childhood trauma, parental substance dependence, parental divorce, and stressful life events have all been cited as environmental risk factors^4,7, 8^. Environmental influences on substance use are typically found to be more pronounced in adolescence and are associated with first use^8-12^. The proportion of variance in alcohol initiation explained by environmental factors has been found to be as high as 76%^13^. Indeed, twin studies have shown that the shared family environment accounts for 23% of the variance in drug use ^14^ but have also highlighted the importance of the environment that is unique to the individual^10^. Typically, environmental influences decrease in importance and genetic effects become more prominent moving from adolescence into early adulthood^9^. The decline in environmental effects is thought in part to reflect the waning influence of authority figures and peers as individuals gain more independence.

The role of the recent shared environment and its effect on adult alcohol and tobacco use is less well studied. Observing the correlations between couples is a useful measure as couples typically share many aspects of the recent environment, whereas their earlier exposures are distinct. Studies have shown that members of a couple have similar levels of substance use^15,16^. The rate of alcohol use disorder in the husbands of female alcoholic probands was found to be 31%^15^ and the correlation for substance dependence symptoms among mothers and fathers of youths in treatment for substance use disorder was found to be 0.4^16^. Furthermore, females who abuse alcohol and nicotine are more likely to have married individuals who do the same^17^.

One explanation for the observed similarities between couple’s substance use is assortative mating. Assortative mating is a pattern of non-random mating whereby individuals with similar phenotypes are more likely to mate with one another and this leads to increased genetic similarity at loci known to be associated with substance use. Although assortative mating has been proposed to increase alcohol dependence correlations between spouses^18^, it may be that indirect genetic effects contribute to phenotypic correlations. Indirect genetic effects occur when the genotype of an individual influences the phenotype of another conspecific individual. For example, people at high genetic risk for alcohol use who drink heavily could create an environment which increases their partner’s risk for alcohol use, such as increased alcohol availability or a stressful environment arising from problematic drinking. In the context of smoking, individuals at low genetic risk for smoking who do not smoke may encourage their partner to quit smoking. Furthermore, as substance use patterns are dynamic, individuals may change their behaviour over time to match that of their partners.

In the present study, we aimed to measure the genetic and environmental contributions to people’s differences in alcohol use and smoking behaviour in a population-based cohort, Generation Scotland: the Scottish Family Health Study (GS)^19,20^. Exploiting the diverse family relationships in GS, we estimate the contribution of shared family, sibling, and couple effects on substance use, and estimate the proportion of phenotypic variance attributable to genetic effects in the presence of these factors. To investigate the potential role of assortative mating, we estimated spousal correlations for alcohol and smoking use phenotypes across 34,987 couple pairs in the UK Biobank (UKB). As assortative mating can lead to increased genetic similarity between individuals, we also estimated the intra-couple polygenic risk score correlations for alcohol and nicotine use phenotypes. We also tested whether a spousal PRS predicted their partner’s phenotype. Finally, we explored whether a functional SNP (rs1229984) in *ADH1B,* that influences alcohol metabolism and is strongly associated with alcohol intake, was associated with levels of the partner’s drinking.

## Methods

### Sample Descriptions

#### Generation Scotland: the Scottish Family Health Study

Generation Scotland: the Scottish Family Health Study (GS) is a family-based cohort recruited via general practitioners across Scotland. Individuals were invited to participate if they were able to recruit at least one other family member aged 18 or over.

##### Genotyping

Genotyping was performed on 20,195 individuals using the Illumina OmniExpress BeadChip. Quality control steps removed individuals with a genotype call rate <98%, SNPs with a call rate of <98%, SNPs with a minor allele frequency less than 1%, or those which deviated from Hardy-Weinberg equilibrium (p < 5 x 10^-6^). Principal component analyses were also performed to remove population outliers^21^. After quality control, 19,904 individuals remained with 561,125 autosomal SNPs.

##### Phenotypes

*Smoking status: S*moking behaviours were assessed as part of a pre-clinical questionnaire. Individuals were asked whether they were current, former or never smokers. Former and current smokers were then collapsed to create an ever/never smoking variable.

*Cigarettes per day:* Participants who had endorsed ever smoking were asked the average amount of cigarettes they smoked daily, currently, or in the past. Cigarettes per day was recorded as an ordinal variable and treated as an interval variable in the present study with 0 = 0, 1 = Less than daily, 2 = 1-4, 3 = 5-9, 4 = 10-14, 5 = 15-19, 6 =20-24, 7 = 25-29, 8 = 30-34, 9 = 35-39, 10 = 40-44, 11 = 45-49, 12 = 50+ cigarettes per day.

*Smoking age of onset:* Lifetime smokers were asked at what age they started smoking and responses were categorized and treated as an interval variable so that 1 = less than 5 years, 2 = 5-9, 3 = 10-14, 4 = 15-19, 5 = 20-24, 6 = 25-29, 7 = 30-34, 8=35-39, 9= 40-44, 10 = 45-49, 11=50+ years of age. Individuals who did not know what age they started smoking were set to NA for age of onset.

*Alcohol consumption:* This was assessed using self-report as part of the pre-clinical questionnaire; participants were asked how many units of alcohol they had consumed in the previous week. A prompt was shown in the questionnaire to provide examples of the typical units of alcohol in each drink type.

*Alcohol misuse:* As part of a GS re-contact study in 2014, 9,618 members of GS completed a follow-up questionnaire as part of the Stratifying Resilience and Depression Longitudinally (STRADL) project^22^. These individuals completed the CAGE questionnaire^23^ which can identify individuals at risk of problem drinking. The CAGE questionnaire consists of 4 questions and provides a total score of 0-4 depending on the number of items endorsed.

The total sample size for each of the GS phenotypes and mean and standard deviations are shown in **Supplementary Table 1**.

**Table 1.** Variance component analysis in Generation Scotland showing the significant environmental and genetic components for each trait when fitted simultaneously.

| **Trait** | **G (Genetic - SNP)**<br>**(S.E.)** | **K (Genetic –**<br>**Kinship) (S.E.)** | **F (Family) (S.E.)** | **C (Couple) (S.E.)** | **S (Sibling) (S.E.)** |
| --- | --- | --- | --- | --- | --- |
| **Units per week** | 0.06 (0.02) | 0.12 (0.06) | 0.07 (0.03) | 0.38 (0.03) | - |
| **CAGE score** | 0.19 (0.03) | - | - | 0.31 (0.04) | - |
| **Smoking status** | 0.22 (0.03) | 0.19 (0.05) | - | 0.29 (0.04) | 0.10 (0.03) |
| **Cigarettes per day** | 0.21 (0.05) | 0.20 (0.06) | - | 0.09 (0.05) | - |
| **Smoking age onset** | 0.14 (0.05) | 0.12 (0.05) | - | 0.09 (0.05) | - |

*Identification of couple pairs:* Using the family and genetic data in GS, couples were identified as those who shared a child. This identified 1,742 genotyped couple pairs.

### UK Biobank

The UK Biobank (UKB) is a prospective population-based sample of 502,629 participants recruited across 22 assessment centers in the United Kingdom from the period of 2006 to 2010^24^. People were invited to participate if they were aged between 40-69 years, were registered with the National Health Service and lived within ∼ 25 miles of an assessment center.

#### Genotyping

Genetic data were available for 487,409 individuals in the UKB and genotyping was performed on either the Affymetrix Axiom array or the UK BiLEVE Axiom array^25^. In order to create a White British unrelated dataset, we removed 131,790 related individuals who were third degree relatives or closer (using a kinship coefficient > 0.044). We identified one individual from each group of relatives by creating a genomic relationship matrix and using a genetic-relatedness cut-off of 0.025 and added these back into the sample (N = 55,745). Quality control steps removed individuals with a genotype call rate <98%, SNPs with a call rate of <98%, SNPs with a minor allele frequency less than 1% or those which deviated from Hardy-Weinberg equilibrium (p < 5 x 10^-6^). After quality control 414,584 autosomal SNPs remained.

#### Phenotypes

*Smoking status:* Smoking status was ascertained as part of a touchscreen interview. Participants were asked whether they were current, previous or never smokers; previous and current smokers were collapsed to make an ‘ever smoker’ phenotype. Former smokers were asked about previous smoking behaviour. Those who endorsed ‘Just tried once or twice’ were classed as never smokers and therefore the phenotype is an ever/never-regular smoker as defined according to the Centre for Disease Control and Prevention (CDC).

*Cigarettes per day:* Current smokers were asked how many cigarettes they smoked on average each day and former smokers about how many cigarettes they previously smoked. If individuals stated they smoked over 150 cigarettes per day this answer was rejected; if they endorsed 100 or over they were asked to confirm this selection.

*Smoking age of onset:* Lifetime smokers were asked using the touchscreen questionnaire how old they were when they first started smoking on most days. Responses under age 5 were rejected and under age 12 were prompted for confirmation.

*Alcohol consumption:* Participants were asked how many of various drink types they normally drank on a monthly and weekly basis and this was converted into a measure of units per week. The full derivation of this measure has been described previously^26^.

*Alcohol misuse:* The AUDIT questionnaire^27^ was administered to a sub-set of the UKB who responded to an online mental health questionnaire follow-up over a one year period in 2017. The AUDIT is a ten-item questionnaire with scores ranging from 0-40 that measures both alcohol consumption (Q1-Q3) and problems with alcohol (Q4-Q10). Three AUDIT scores were created based on the score for all questions (AUDIT-T), on the questions measuring alcohol consumption and frequency (AUDIT-C [Q1-Q3]), and on those measuring problems with alcohol or alcohol abuse (AUDIT-P [Q4-Q10]). These measures have been described in greater detail previously^28^.

The total sample size for each of the UKB phenotypes used in the present study and mean and standard deviations are shown in **Supplementary Table 1**.

### Identification of couple pairs

Participants were assigned to couple pairs on the basis of a shared household identifier. Individuals who shared a household, reported living in a household with 2 individuals, and who reported living with a husband, wife, or partner were selected. Any couples with an age gap of greater than 10 years were removed, as were couples whose parental ages matched for either parent. After further selecting White British unrelated individuals from this group there were 34,987 opposite-sex pairs available for analysis. 407 same-sex couples were also identified using the above algorithm. Due to the lower number of same-sex pairs genetic correlations were not analysed in these individuals although phenotypic correlations were estimated.

### Heritability analyses in Generation Scotland

Genetic and environmental effects were estimated in GCTA v1.91 using linear mixed models ^29^ by fitting a pedigree kinship matrix and a SNP matrix (genetic relationship matrix) alongside 3 matrices representing the environment shared by nuclear families (parents and children) (F), couples (identified by a shared child) (C), and siblings (S).

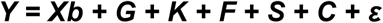

Y is a vector representing the substance use trait of interest and b is the effect of X, a matrix of values that represents the fixed effect covariates of age, sex, and 20 principal components. The genetic effects are represented by G (SNP matrix) and K (pedigree kinship matrix), the three environmental components are F, S, and C, and *ε* the residual error term. This method was first described by Xia et al. ^30^, and the construction of the genetic and environmental matrices are described in more detail in the **Supplementary Material.** Briefly, the G component captures variance explained by common SNPs, the K component captures additional genetic effects by modelling pedigree relationships (achieved by setting all entries in the SNP matrix less than 0.025 to 0). The F component represents nuclear family members by setting the relationship matrix coefficient to 1 if individuals were parent-offspring, sibling or couples. Similarly, the S component represents sibling pairs and the C component couple pairs.

The most parsimonious model was selected by performing backward stepwise selection. The initial model included all 5 components (GKFCS) and components were removed iteratively if they failed to meet significance in the likelihood ratio test (LRT) and Wald tests (α=5%) and among the components satisfying this condition it had the highest (least significant) P value in the Wald test. This process was repeated until all the remaining components were significant in either the LRT or Wald test. The population prevalence for smoking status was 48% and used to convert the estimates for this trait from the observed scale to the liability scale.

### Polygenic risk scores

Polygenic risk scores (PRS) were created using publicly available genome-wide association studies (GWAS) of the substance use phenotypes analysed in this study. For alcohol consumption this was the GWAS of alcohol consumption in the first release of the UKB (N=112,117)^26^. This GWAS contained no GS individuals and we removed any UKB couples who were present in this GWAS from the current dataset when performing PRS analyses. For the smoking phenotypes we used the Tobacco and Genetics Consortium GWAS of smoking status (N=69,409), cigarettes per day (N=38,181), and age of onset (N=22,438)^31^.

PRS were created in PRCise-2^32^ using raw QC’d genotype data and a MAF cut-off of 0.01. The parameters of r^2^=0.1 and window=250 kb were used to create independent SNPs and the scores for p-value thresholds from 0.00005-0.5 created in increments of 0.00005. The score which explained the most variance in the trait of interest was then used for downstream analyses (**Supplementary Figures 1-4**). PRS were regressed onto the first 4 principal components to correct for population stratification and the residuals taken for analyses.

**Figure 1.**
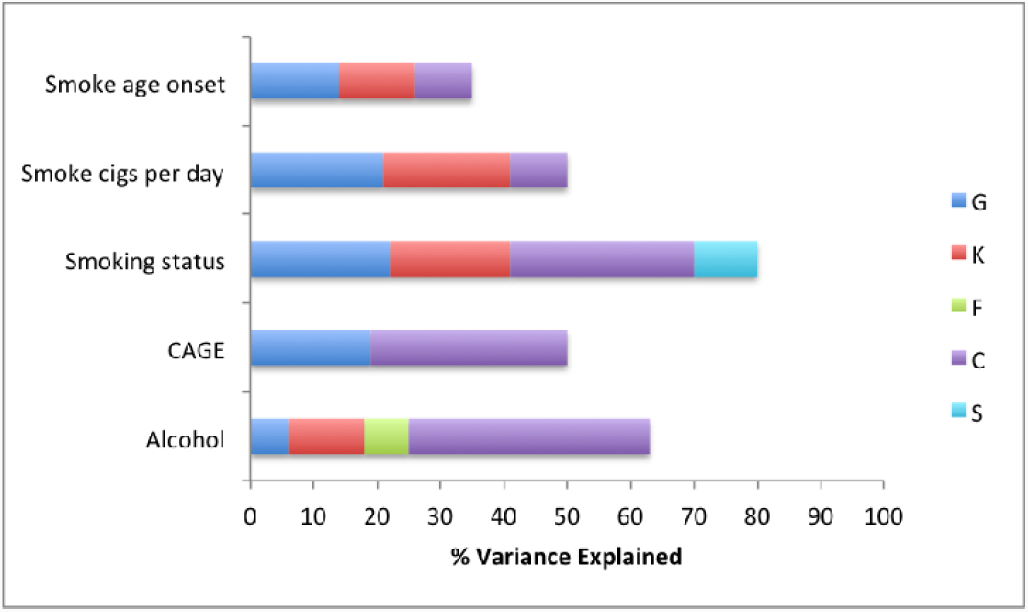
Proportion of variance in substance use traits explained by genetic and environmental components in Generation Scotland. G = genetic, K = kinship, F = nuclear family, C = couple, S = sibling.

### Statistical analyses

Phenotypic associations were tested in R using linear regression models. Phenotypes were regressed onto age and sex, and then age, sex and test-center (categorical) and the residuals from these used for regression analyses. Association between PRS (residualized for principal components) were also performed in R using linear models. Variables were scaled to have a mean of 0 and a standard deviation of 1 and therefore the reported beta are standardized. Permutation tests were carried out to test the independence of couple phenotypes using the *coin* package in R and 10,000 Monte Carlo re-samplings^33^.

## Results

The total sample size for each of the substance abuse phenotypes available in Generation Scotland and UK Biobank is shown in **Supplementary Table 1**, along with the mean and standard deviations for each trait. The variance explained by each genetic and environmental component in the Generation Scotland cohort is shown in **Table 1** and **Figure 1**. Significant genetic effects were detected for all traits. Units per week of alcohol has the lowest SNP-based genetic estimate, accounting for 6% of the variance (S.E.=0.02). The CAGE score’s SNP-based genetic contribution was 19% (S.E.=0.03). Smoking status had the highest estimate (G=0.22, S.E.=0.03). With the exception of CAGE score, the kinship component significantly contributed to the variance explained for all traits, with values between 12% and 20%. This suggests a role for additional genetic effects such as rare variants or epistatic effects that are detectable when analysing close relatives. The sum of the G and K components is comparable to narrow-sense heritability estimates^30^ and therefore the total genetic contribution to units per week and CAGE score was 18% and 19% respectively. For smoking status, smoking age of onset and cigarettes per day the narrow sense heritability estimates were 41%, 26% and 41% respectively.

The most significant environmental contribution across all traits was the couple component (C). The contribution was 29% (S.E.=0.04) for smoking status and 9% (S.E. = 0.05) for cigarettes per day. The largest contribution, of 38% (S.E.=0.03), was to the phenotypic variance explained for units of alcohol consumed per week. Modest early environmental effects were also detected for units per week and smoking status, as follows: the nuclear family component (F) explained 7% of the variance (S.E.=0.03) in units per week and the shared sibling environment (S) explained 10% of the variance in smoking status (S.E.=0.03) (**Table 1**) (**Figure 1**).

The results of the full backward stepwise model selections are shown in **Supplementary Table 2**. For smoking status, 80% of the variance was explained suggesting that only 20% of the variance can be apportioned to other environmental effects or sampling error (**Figure 1**). The total variance explained in units per week was 63% and for cigarettes per day and CAGE score the total variance explained was 50%. For some traits the majority of phenotypic variance was unexplained: only 35% of the variance in age of smoking onset was explained, suggesting that the majority of the variance in this trait is influenced by unique environmental factors or shared factors that are not captured by the current model. The unexplained variance could also be attributed to measurement error. Substance use can be difficult to measure as it relies on accurate recall of behaviours which change across the lifespan.

**Table 2.** Phenotypic correlations between substance use phenotypes in opposite-sex couples in UKB (N=34,987 pairs). N shown in trait column reflect the N where both members of the couple had available phenotype data

|  | Baseline |  |  | Age + Sex |  |  | Age + Sex + Test Centre |  |  |
| --- | --- | --- | --- | --- | --- | --- | --- | --- | --- |
| Trait | Beta | SE | P | Beta | SE | P | Beta | SE | P |
| **Smoking Status**<br>**N=34,732** | 0.22 | 0.005 | $< 2 \times 10^{-16}$ | 0.21 | 0.005 | $< 2 \times 10^{-16}$ | 0.21 | 0.005 | $< 2 \times 10^{-16}$ |
| **Smoking Age Onset**<br>**N=5226** | 0.09 | 0.013 | $9 \times 10^{-12}$ | 0.09 | 0.01 | $3 \times 10^{-12}$ | 0.09 | 0.01 | $1 \times 10^{-10}$ |
| **Smoking Cigs per Day**<br>**N=4749** | 0.13 | 0.01 | $< 2 \times 10^{-16}$ | 0.13 | 0.01 | $< 2 \times 10^{-16}$ | 0.12 | 0.01 | $< 2 \times 10^{-16}$ |
| **Alcohol units per week**<br>**N=31,263** | 0.52 | 0.005 | $< 2 \times 10^{-16}$ | 0.52 | 0.005 | $< 2 \times 10^{-16}$ | 0.52 | 0.005 | $< 2 \times 10^{-16}$ |
| **AUDIT score**<br>**N=6787** | 0.48 | 0.011 | $< 2 \times 10^{-16}$ | 0.47 | 0.01 | $< 2 \times 10^{-16}$ | 0.47 | 0.01 | $< 2 \times 10^{-16}$ |
| **AUDIT-C**<br>**N=6787** | 0.51 | 0.011 | $< 2 \times 10^{-16}$ | 0.50 | 0.01 | $< 2 \times 10^{-16}$ | 0.50 | 0.01 | $< 2 \times 10^{-16}$ |
| **AUDIT-P**<br>**N=6787** | 0.12 | 0.009 | $< 2 \times 10^{-16}$ | 0.11 | 0.009 | $< 2 \times 10^{-16}$ | 0.11 | 0.009 | $< 2 \times 10^{-16}$ |

All traits in GS showed a significant amount of variance explained by the couple environment (C, **Table 1**). The within-couple phenotypic associations in GS are shown in **Supplementary Table 3**. Smoking status (b=0.19 (S.E.=0.02)), alcohol consumption (b=0.26 (S.E.=0.03)), and CAGE score (b=0.22 (S.E.=0.05)) were all significantly associated within couple pairs. After residualizing the traits for age, sex and recruitment area a nominally significant couple association was also observed for cigarettes per day (b=0.10, (S.E.=0.05)) (**Supplementary Table 3**) and the couple association for alcohol consumption became stronger (b=0.41, (S.E.=0.02)).

**Table 3.** Association between couple polygenic risk scores (PRS) in UKB for substance use traits.

|  | **Association between couples PRS UKB** |  |
| --- | --- | --- |
| **PRS trait** | **PRS Beta (S.E.)** | **P-Value** |
| **Alcohol Consumption** | 0.007 (0.008) | 0.35 |
| **Smoking Status** | 0.013 (0.005) | 0.02 |
| **Cigarettes per day** | 0.007 (0.005) | 0.18 |
| **Age smoking onset** | 0.006 (0.005) | 0.29 |

Similar phenotypic associations were observed between members of couple pairs in the UKB. Smoking status, cigarettes per day, age of smoking onset, units of alcohol per week, and AUDIT scores were all significantly associated within couple pairs (**Table 2**). Controlling for age, sex and recruitment center did not significantly alter the observed associations. AUDIT scores were strongly associated between members of a couple (b=0.48 (S.E.=0.01), p < 2 x 10^-16^) as were units per week (b=0.52 (S.E.=0.005), p < 2 x 10^-16^). Smoking status, cigarettes per day and age of smoking onset were more modestly associated within couples (b=0.09-0.22, p < 9 x 10^-12^).

There were 407 same-sex pairs identified in the UKB and the phenotypic correlations between these individuals are presented in **Supplementary Table 4**. Although there were very few individuals for some phenotypes, the correlations for the phenotypes with larger sample sizes (smoking status, units per week) appear similar to those observed in opposite-sex pairs.

**Table 4.** Association between male PRS and female PRS and partner and own phenotype in the UKB

Table 3) Association between couple polygenic risk scores (PRS) in UKB for substance use traits.
| PRS association<br>(standardized $\beta$, standard error, p-value and $r^2$) | |
| --- | --- |
| Males –PRS with partner phenotype |  |
| Alcohol PRS | $\beta = 0.035$ (S.E. = 0.007) $p = 2.3 \times 10^{-7}$, $r^2=0.11\%$ |
| Age smoking onset PRS | $\beta = -0.008$ (S.E.= 0.01) $p = 0.47$, $r^2=0\%$ |
| Smoking status PRS | $\beta = 0.014$ (S.E. = 0.005) $p= 0.01$, $r^2=0.02\%$ |
| Cigarettes per day PRS | $\beta = -0.012$ (S.E. = 0.01) $p = 0.25$, $r^2=0.004\%$ |
| Females –PRS with partner phenotype |  |
| Alcohol PRS | $\beta = 0.041$ (S.E. = 0.007) $p = 3.1 \times 10^{-9}$, $r^2=0.15\%$ |
| Age smoking onset PRS | $\beta = 0.011$ (S.E. = 0.008) $p = 0.17$, $r^2=0.006\%$ |
| Smoking status PRS | $\beta = 0.013$ (S.E. = 0.005) $p= 0.02$, $r^2=0.01\%$ |
| Cigarettes per day PRS | $\beta = 0.006$ (S.E. = 0.009) $p = 0.48$, $r^2=0\%$ |
| Own –PRS with own phenotype |  |
| Alcohol PRS | $\beta = 0.078$ (S.E. = 0.004) $p < 2 \times 10^{-16}$, $r^2=0.62\%$ |
| Age smoking onset PRS | $\beta = 0.020$ (S.E. = 0.007) $p = 0.002$, $r^2=0.04\%$ |
| Smoking status PRS | $\beta = 0.043$ (S.E. = 0.004) $p < 2 \times 10^{-16}$, $r^2=0.18\%$ |
| Cigarettes per day PRS | $\beta = 0.034$ (S.E. = 0.007) $p < 9 \times 10^{-7}$, $r^2=0.11\%$ |

Couple correlations can arise because of assortative mating, whereby individuals with similar phenotypes mate, potentially resulting in greater genetic similarity between members of a couple. In order to test this, polygenic risk scores (PRS) were created for each substance use trait using GWAS summary statistics from independent samples. The associations between couples PRS were then tested in the UKB as there were more couple pairs available (N=34,987 vs N=1742 in GS). In the UKB a weak but significant correlation between partners smoking status PRS was observed (b=0.013, S.E.=0.005, p=0.02) (permutation p-value = 0.006). No significant correlations were observed between partners’ alcohol consumption PRS in the UKB sample (**Table 3**).

Individuals’ PRSs were tested for association with the partner’s substance use phenotypes. Male alcohol consumption PRS was significantly positively associated with female partner’s alcohol consumption in the UKB (b=0.035, S.E.=0.007, p=2 x 10^-7^, r^2^=0.11%) (permutation p-value < 2 x 10^-16^). The same was observed for female alcohol consumption PRS – a significant association with male partner phenotype was found (b=0.040, S.E.=0.007, p=3 x 10^-9^, r^2^=0.15%) (permutation p-value < 2 x 10^-16^) (**Table 4)**. The association between alcohol consumption PRS and partner consumption is weaker, and explains less of the variance than the association with an individual’s own alcohol consumption (b=0.078, S.E.=0.004, p < 2 x 10^-16^, r^2^=0.62%) (**Table 4**). Significant associations between male and female smoking status PRS and partner phenotype were also observed in the UKB. The PRSs for age of smoking onset and cigarettes per day were not associated with corresponding partner phenotypes in UKB (**Table 4**).

The association between the rs1229984 *ADH1B* SNP and units per week was also tested in the UKB. rs1229984 is a non-synonymous SNP in the alcohol dehydrogenase 1B gene (*ADH1B);* the minor allele (A) carriers have a version of ADH1B that oxidizes alcohol more rapidly, and as such A carriers are at a reduced risk for alcohol use disorder ^34,35^. The minor allele frequency of rs1229984 in the UKB couples sample was 2.6% (A allele). In the present sample of UKB individuals, those with the AA/AG genotype at rs1229984 drank 4.9 units per week on average (S.E.=0.08) compared to GG individuals who drank 6.276 units (S.E.=0.02) (p< 2 x 10^-16^). Units per week of alcohol for each genotype in the UKB couples sample is shown in **Supplementary Figure 5**.

We next took all the individuals with a GG genotype and split them according to whether they had a partner with the GG genotype (GG-G) or an A carrier partner (AG or AA) (GG-A). GG-G individuals consumed on average 6.301 units per week (S.E.=0.02). GG-A individuals consumed significantly less than their GG-G equivalents, 5.8 units per week (S.E.=0.10) (p < 9 x 10^-7^). Having a partner who carries an rs1229984 A allele was, therefore, associated with a mean of 0.5 units less alcohol per week intake amongst G carrier individuals.

## Discussion

In the present study, using genotyping and family relationships data, we show that there are significant genetic and environmental contributions to substance use in a general population sample, Generation Scotland. The effect of the shared couple environment was particularly pronounced and contributed significantly to the variance in each trait. In support of this, we report significant phenotypic correlations within couples for all of the substance use traits in the UK Biobank. In order to test whether this was due to assortative mating we analysed the correlations between partners’ substance use polygenic risk scores (PRS) in the UKB. Whereas there was no significant association between alcohol consumption PRS within couples, an individual’s alcohol consumption PRS associated with their partner phenotype. Furthermore, the presence of the rs1229984 A allele in a partner was associated with reduced alcohol intake in individuals with GG genotypes at this locus. For smoking phenotypes, there was some evidence that PRS for smoking status were associated and PRS also tended to associate with partner phenotype.

The narrow-sense heritability of alcohol use phenotypes reported in this study (sum of G and K) are lower than those generally reported in the literature. The narrow-sense heritability of alcohol consumption and CAGE score was estimated to be 18% and 19% respectively. Broad sense heritability estimates of 25-61% for alcohol consumption have previously been reported from studies of twins^1,2,5^. The SNP effects for alcohol consumption were estimated at 6%, which again are lower than, but closer to, estimates reported in the UK Biobank for alcohol consumption (13%) and AUDIT scores in the 23andMe sample (12%)^28,36^. Previous studies have suggested that genetic interactions and improper modeling of the environment can inflate heritability estimates^37, 38^. The narrow-sense heritability estimates for cigarette smoking were higher ranging from 26% for age of onset to 41% for smoking status and cigarettes per day. These are somewhat lower than heritability estimates from twin studies (typically 45-80%). We report, the SNP heritability of smoking status (22%), age of smoking onset (14%) and cigarettes per day (21%). The SNP heritability of smoking status has previously been estimated at 17% in the UKB, similar to our estimate in GS^39^.

Early environmental factors, such as those shared by families and siblings, did not appear to explain large amounts of variance in adult substance use in this sample. Shared family environment was estimated to explain 7% of the variance in units per week of alcohol consumed. Given that the age range of the Generation Scotland sample is 18-99 years, it is likely that most members of a nuclear family no longer share a household and so the family component should represent early shared environment in this study. Parental expectations, attitudes and alcohol use have all been shown to influence adolescent alcohol use. Fewer studies have examined the effect of familial influences into adulthood; however, a family history of alcohol abuse and^40^ age of first drink^41^ are associated with alcohol abuse in later life, although it is unclear whether these represent genetic or environmental factors. A family study from the Netherlands found non-shared environmental factors to explain the majority of the variance in alcohol consumption and found no evidence of cultural transmission influencing adult alcohol use^42^. The findings from our study suggest that there may be a small contribution of family environment on drinking patterns in later life; these discrepant findings may be due to cultural differences between the samples.

A significant shared sibling effect was detected for smoking status, explaining 10% of the variance in this trait. Sibling effects can represent genetic or environmental effects; however, as we model genetic effects simultaneously in our model the component captures the effect of the early shared environment. Previous studies have shown sibling concordance in smoking status and this is greater when siblings report a high degree of social connectedness^43^. This suggests we may be detecting shared peer effects or the influence of one sibling’s smoking status on the other^44^.

The proportion of phenotypic variance in alcohol consumption explained by the couple environment in GS was substantial at 38%. The phenotypic correlations for all alcohol use phenotypes in both GS and the UKB were high. The correlations between partner alcohol consumption, AUDIT score and AUDIT-C (consumption) were 0.47-0.52 in the UKB; however, the AUDIT-P (problems) correlation was smaller (0.12). Similarly, in GS the alcohol consumption correlation in GS was high (0.4) whereas the correlation between partner CAGE score was lower (0.22-0.24), demonstrating that alcohol consumption is more strongly correlated between partners than patterns of alcohol abuse in these samples. Correlations between partners can be driven by assortative mating. Partner alcohol consumption PRS were not correlated in the UKB; however, alcohol consumption PRS did predict partner phenotype in the UKB for both males and females. Also, having a partner who carries an A allele at the rs1229984 locus was associated with lower alcohol intake among G carriers of this SNP. PRS typically explain very little of the variance in the traits they predict (<1%) and therefore the lack of correlation between couples PRS does not rule out assortative mating as an explanation for the couple similarities.

Indirect genetic effects occur when the genotype of one individual influences the phenotype of another. The influence of genotype on partners’ substance use may be via the contribution of that genotype to the environment, such as creating high or low exposure to alcohol. This is similar to the genetic nurture effect described by Kong et al^45^. Using PRS for educational attainment, they show that the offspring of parents with higher PRS have greater educational attainment themselves, even when they do not inherit the ‘education-associated’ alleles. The nurturing environment provided by the parents with higher PRS is proposed to increase educational attainment of the offspring. In the case of alcohol consumption, partner genotype may lead to higher or lower alcohol exposure, or different attitudes towards alcohol use, which could lead to changes in partner substance use. It is difficult to distinguish between indirect genetic effects and assortative mating from our results alone, and it is possible that both are occurring. Furthermore, levels of alcohol consumption between members of a couple may become more similar over time, potentially in response to shared environmental factors such as life stress or social deprivation. Longitudinal samples or samples with more couple pairs are required to tease apart the potential contributions of each of these factors to couple substance use behaviour.

For the smoking phenotypes the variance explained by the couple environment ranged from 9-29%. As age of smoking onset and smoking status are typically determined during adolescence or early adulthood, behaviour convergence is less likely to explain the couple correlations observed for these traits. Assortative mating may explain some of this effect as weak but significant correlations between couples PRS were observed for smoking status in the UKB. No significant couple correlations were observed for age of smoking onset or cigarettes per day; however, it should be noted that the PRS for smoking initiation only weakly predicted age of onset in the UKB (**Table 4**) and therefore may be a poor instrument to test for assortative mating. Assortative mating can also be measured by assessing the gametic phase disequilibrium (GPD) of trait increasing alleles across the genome ^46^. GPD, as a consequence of assortative mating, manifests as an increased likelihood of carrying trait-increasing alleles across the genome, independent of linkage disequilibrium. Deriving PRS from odd numbered chromosomes and analyzing the correlation with PRS derived from even numbered chromosomes can quantify GPD. A recent study, which also used UKB individuals, found no evidence of GPD for alcohol use or smoking behaviour providing additional evidence that assortative mating does not significantly contribute to the phenotypic couple correlations reported in the literature^46^.

There are a number of limitations to this study. We used PRS to assess the contribution of assortative mating to phenotypic couple correlations and found some evidence of this for smoking status. The presence of assortative mating therefore has implications for the heritability estimates of smoking status. By incorrectly modeling the couple effect as an environmental effect we reduce the residual error term in the model and may inflate the heritability estimates; however, in the absence of longitudinal data it is difficult to determine whether assortative mating or shared couple environment is responsible for the correlation between substance use phenotypes. Another limitation is that the substance use phenotypes are based on self-report, and for the initiation of smoking and cigarettes per day, rely on retrospective accounts which can be unreliable. Also, the definition of a never smoker according to the CDC is someone who has smoked less than 100 cigarettes in their lifetime. We were able to create a phenotype similar to this in the UKB, but for GS we had to dichotomize smokers into never vs ever smokers and therefore these phenotypes are not directly comparable. Finally, assigning individuals to couples was done differently in GS and UKB. Genetic data was used to identify couples who shared a child in GS, but it is possible that these individuals did not share a household at the time of recruitment. Given that GS was recruited through family participation this is less likely but cannot be ruled out. Similarly for UKB, couple data was not linked in the database but using strict exclusion criteria we were able to generate couples from the household data provided. It is more likely that we excluded potential couples from the UKB.

In conclusion, we find that the similarities within couples explains a large amount of the variance in substance use phenotypes, particularly for alcohol consumption. It is unclear whether this is due to shared environmental factors, assortative mating or indirect genetic effects. Future studies analyzing the contribution of couple effects to substance use would benefit from using longitudinal data to better understand how behaviours change as individuals enter relationships and larger family samples with more couple pairs are needed to model the effect of couple’s genotype alongside an individual’s own genetic effects. It is important to understand the effect of assortative mating on substance use as it increases the likelihood of children inheriting any genetic risk for substance use disorders alongside the additional impact of an adverse early environment from two parents with substance use problems^18^. If substance use behaviours converge to cause spousal similarities then this is a potential modifiable risk factor to consider when addressing substance abuse as targeted interventions can be developed for vulnerable individuals. Given the magnitude of the couple correlations reported here, it might be worthwhile to consider the substance use of someone’s partner when any interventions to reduce intake are implemented.

## Acknowledgements

Generation Scotland received core support from the Chief Scientist Office of the Scottish Government Health Directorates [CZD/16/6] and the Scottish Funding Council [HR03006]. Genotyping of the GS:SFHS samples was carried out by the Genetics Core Laboratory at the Wellcome Trust Clinical Research Facility, Edinburgh, Scotland and was funded by the Medical Research Council UK and the Wellcome Trust (Wellcome Trust Strategic Award “STratifying Resilience and Depression Longitudinally” (STRADL) Reference 104036/Z/14/Z). This research was conducted using the UK Biobank Resource, application number 4844, and was supported by the Centre for Cognitive Ageing and Cognitive Epidemiology (CCACE), which was funded by the Medical Research Council and the Biotechnology and Biological Sciences Research Council (reference MR/K026992/1). AJM, DP and IJD received support from an MRC Mental Health Data Pathfinder Grant (reference MC_PC_17209). IJD received support from the MRC CCACE grant (reference MR/K026992/1).

## Author contributions

AC, AM, BHS, CH, DP, GD, MJA, SP contributed to the data acquisition, quality control and processing of the samples for this study. Manipulation of genetic data and quality control was performed by TKC, CH, MJA, DMH, CX and GD. TKC and CX were responsible for the study concept and design. Data analysis and interpretation of findings were performed by TKC, CX and AMM. The manuscript was drafted by TKC. IJD, AMM and CX provided critical revision of the manuscript for important intellectual content. All authors critically approved the content of the manuscript and approved the final version for publication.

