## Supplementary Materials for "Genetic and shared couple environmental contributions to smoking and alcohol use in the UK population"

G: Genomic relationship matrix: The GRM for the GS individuals was calculated in GCTA using the following formula:

$Ajk=\frac{1}{N}\sum_{i=1}^{N} \frac{(xij-2pi)(xik-2pi)}{2pi(1-pi)}$(Yang *et al.*, 2011)

where i is a SNP, x is the allele count of the minor allele for individual j or k at each SNP (i). pi is the minor allele frequency of SNP i and N is the total number of SNPs. G ~ *N* (0, GRM_g_$\boldsymbol{\sigma}_{\boldsymbol{g}}^{\boldsymbol{2}}$).

K: Kinship relationship matrix: The K matrix was created by modifying the values of G. Relationship co-efficients that were less than or equal to 0.05 were set to 0. The threshold of 0.05 was applied as it separates closely and distantly related individuals (Zaitlen et al., 2013). K ~ *N* (0, GRM_kin_$\boldsymbol{\sigma}_{\boldsymbol{kin}}^{\boldsymbol{2}}$).

C,S,F: Environment relationship matrices (ERM) were created to represent shared environmental effects from the different family relationships in GS. Each ERM was created by making an N x N matrix and all entries set to 0 with the diagonal entries set to 1. Off diagonal entries were 1 if 2 families shared the environment of interest (couple, spouse etc). C represents the shared couple environment, S represents the shared sibling environment, and F represents the shared nucular family environment of individuals who once likely shared the same household.

Supplementary Figures

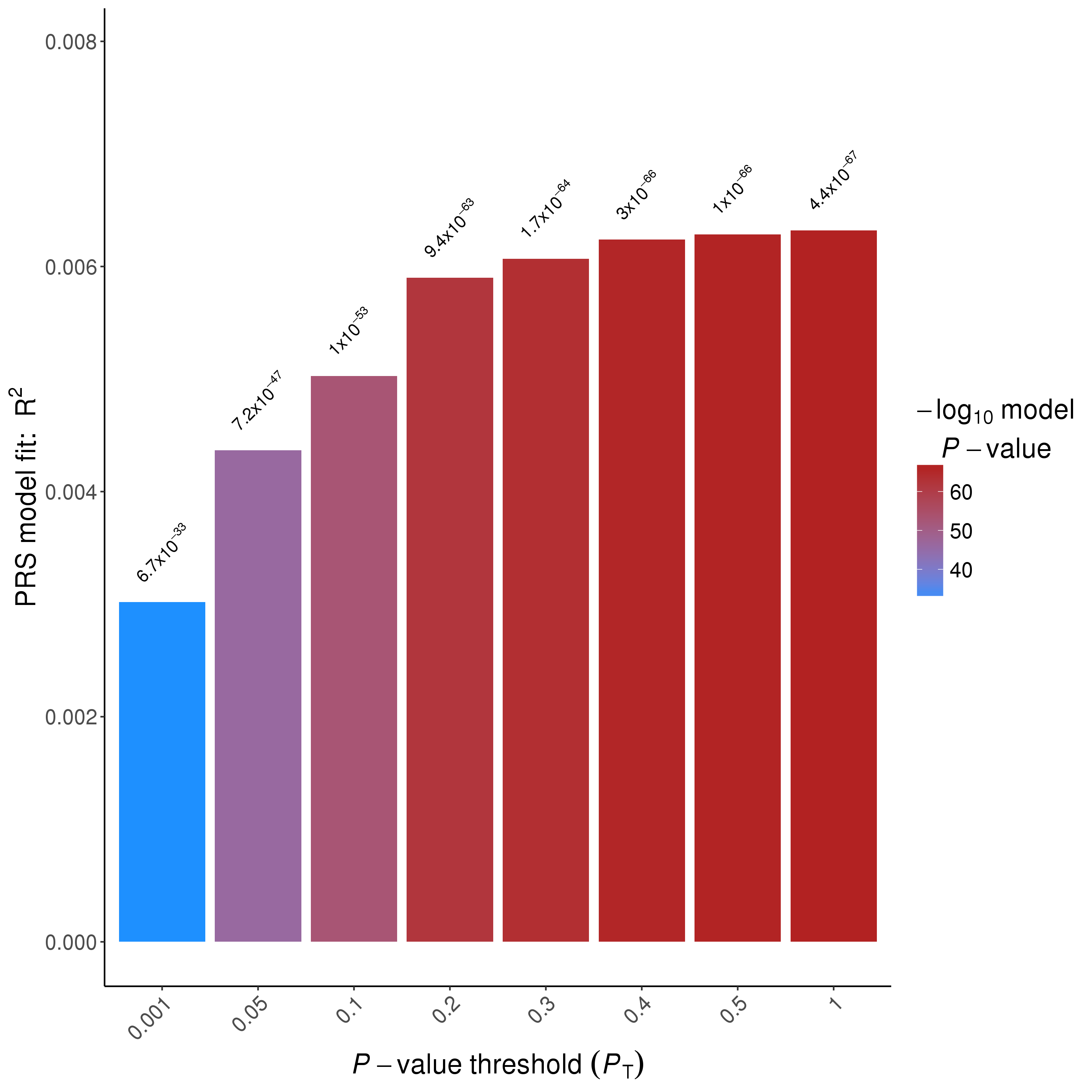

SF1: Best PRS p-value threshold for alcohol consumption in UKB estimated at p < 1. P-value for association between trait and PRS shown above bars.

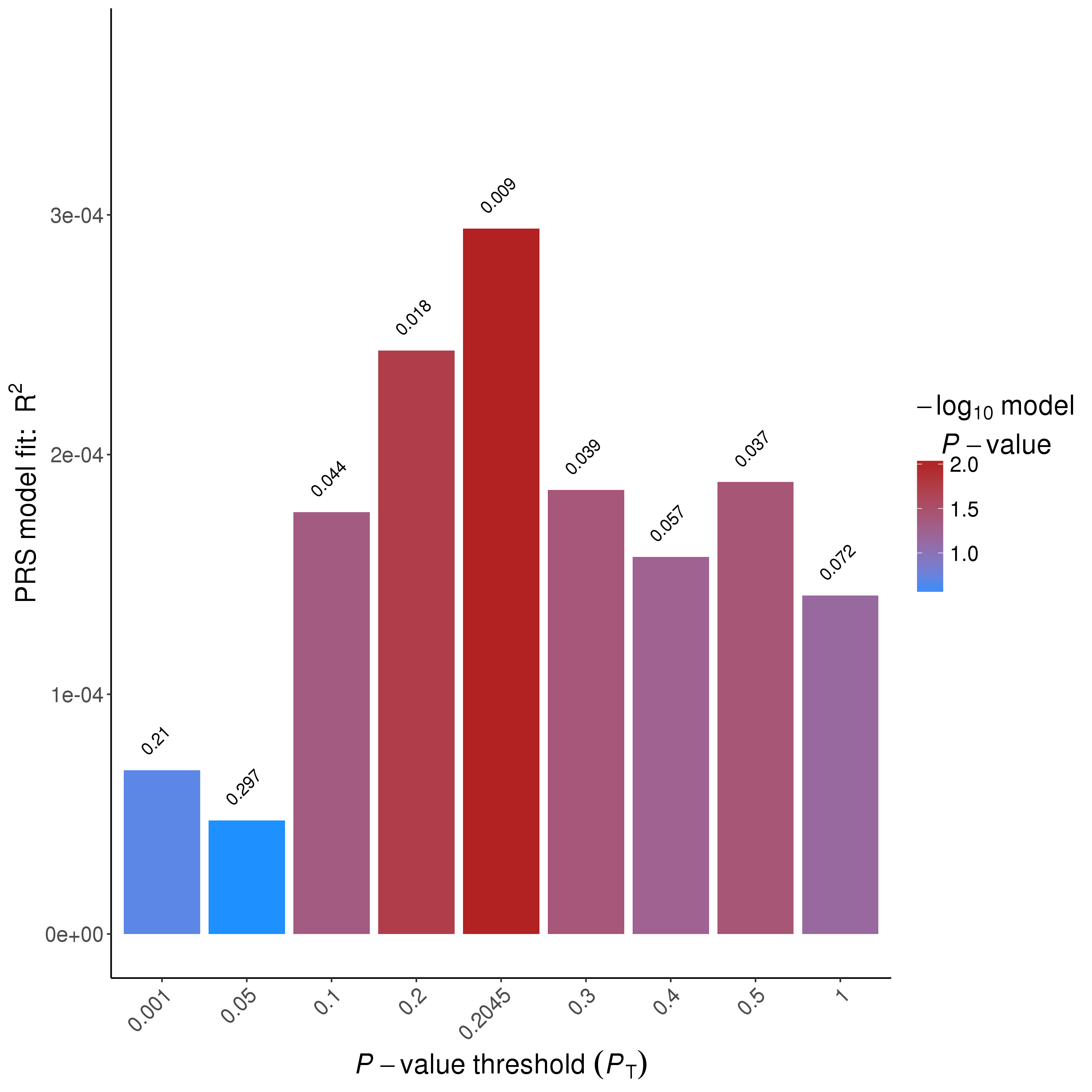

SF2: Best PRS p-value threshold for smoking age of onset in UKB estimated at p ≤ 0.2045. P-value for association between trait and PRS shown above bars.

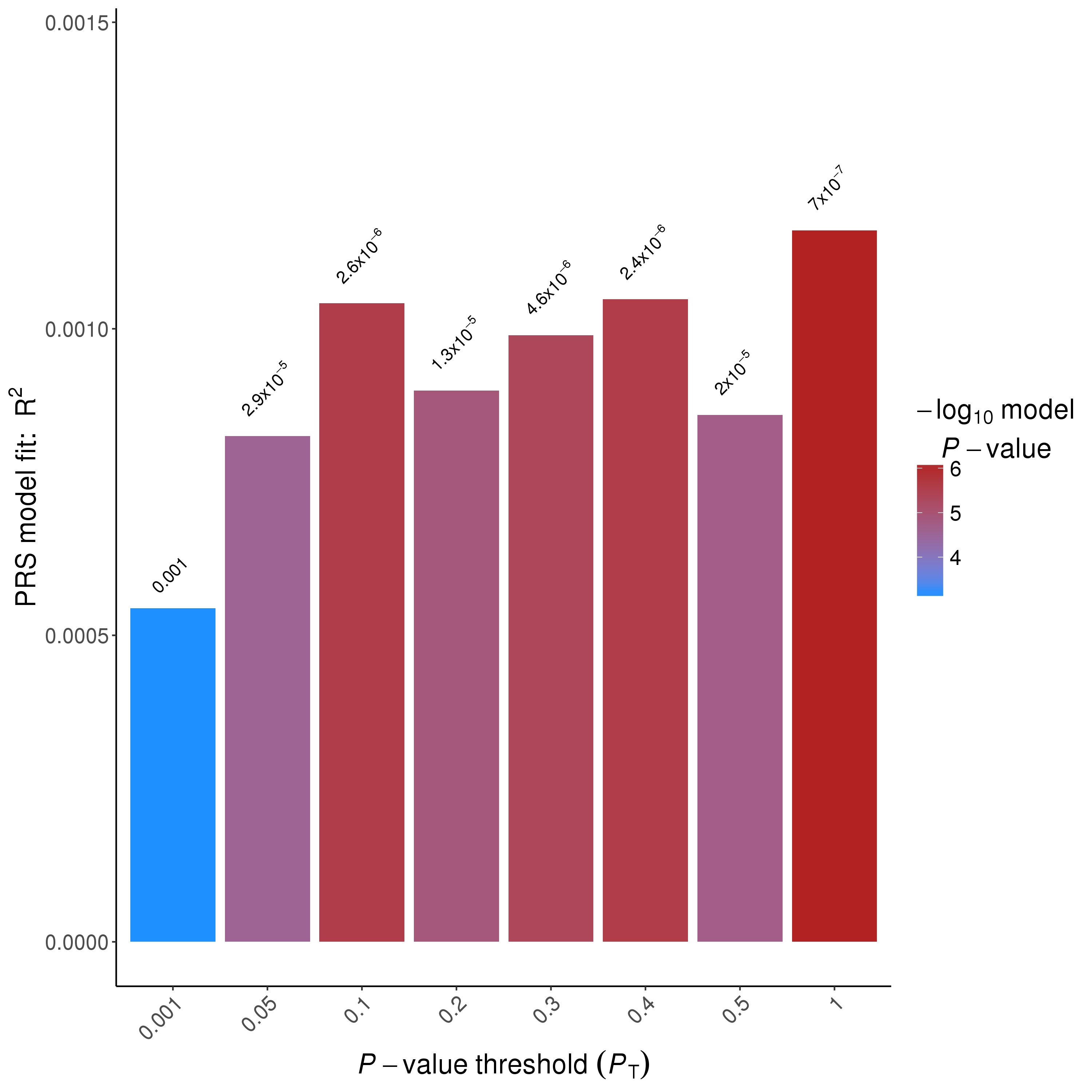

SF3: Best PRS p-value threshold for cigarettes per day in UKB estimated at p ≤ 1. P-value for association between trait and PRS shown above bars.

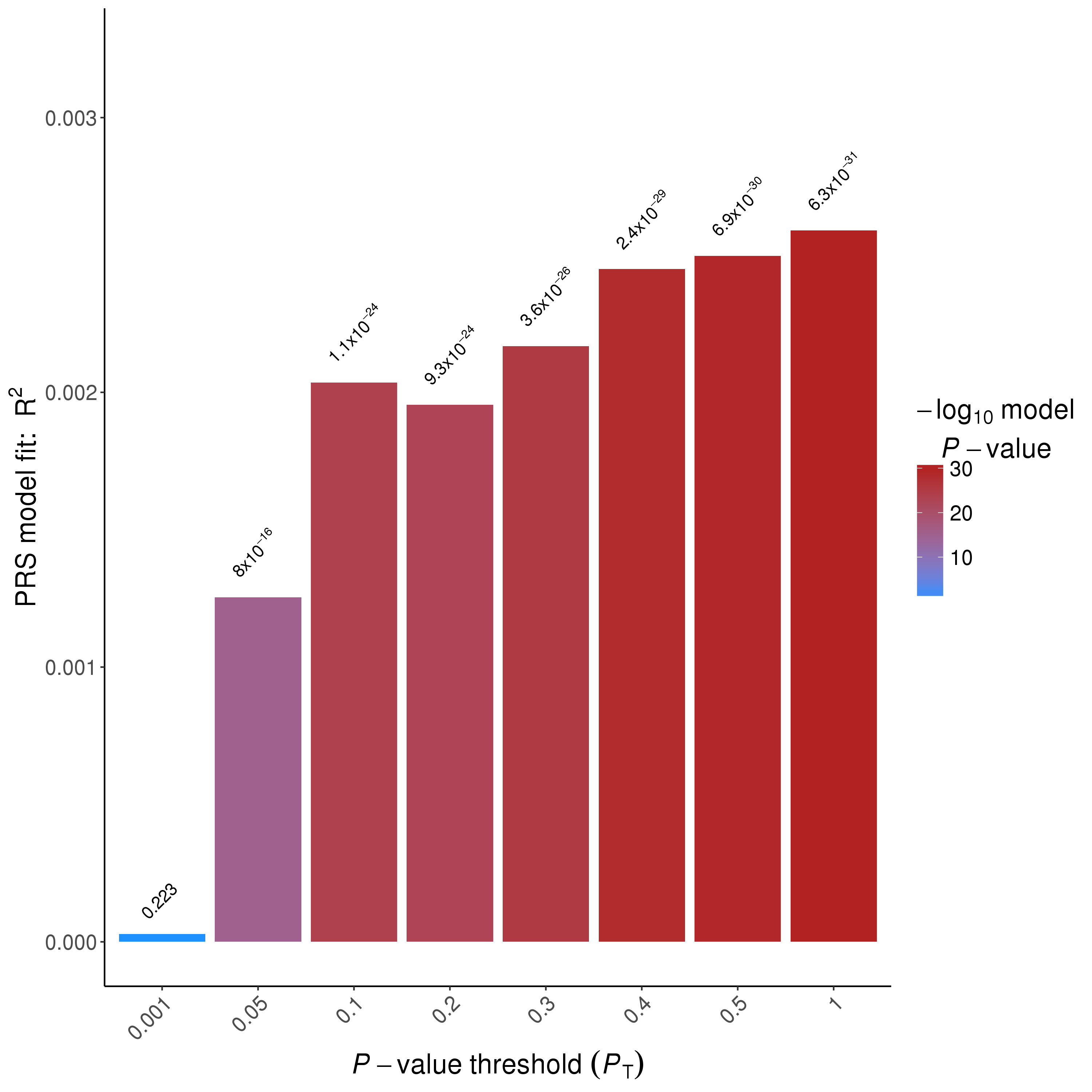

SF4: Best PRS p-value threshold for ever smoking in UKB estimated at p ≤ 1. P-value for association between trait and PRS shown above bars.

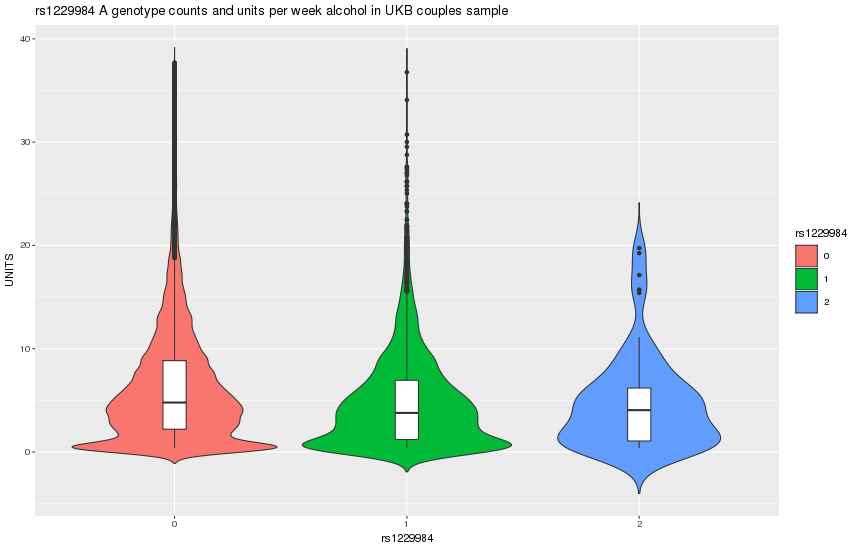

**SF5**: Weekly alcohol intake in units in UKB according to rs1229984 genotype.
