## Supplementary Tables for "Genetic and shared couple environmental contributions to smoking and alcohol use in the UK population"

| **Generation Scotland** | | | **UK Biobank** | | |
| --- | --- | --- | --- | --- | --- |
| **Smoking behaviour** | | | | | |
|  | **Mean** | **S.D.** |  | **Mean** | **S.D.** |
| **Smoking status**  **(N=19,377)** | 47.3% ever smoked | | **Smoking status**  **(N=69,714)** | (46.9% ever regularly smoked) | |
| **Cigarettes per day**  **(N=8461)** | 4.2  (Category 4 corresponding to 10-14 cpd) | 2.2 | **Cigarettes per day**  **(N=21,374)** | 18.7 | 10.1 |
| **Age of smoking onset**  **(N=9152)** | 4.0  (Category 4 corresponding to 15-19 years) | 0.9 | **Age of smoking onset**  **(N=23,199)** | 17.3 | 3.8 |
| **Alcohol Use** | | | | | |
| **Units per week**  **(N=17,546)** | 10.9 | 11.5 | **Units per week**  **(N=65,722)** | 15.9 | 15.4 |
| **CAGE Score**  **(N=7326)** | 0.6 | 0.9 | **AUDIT-T score**  **(N=22,853)** | 4.7 | 3.8 |
|  |  |  | **AUDIT-C score**  **(N=22,853)** | 4.2 | 2.7 |
|  |  |  | **AUDIT-P score**  **(N=22,853)** | 0.6 | 1.7 |

ST1: Total sample size, mean and standard deviations for the Generation Scotland sample and the UK Biobank. The descriptive statistics for the total GS sample are presented whereas only those individuals contribution to the couple-pair analysis in UKB are shown.

| **Units per week** | | | | | |
| --- | --- | --- | --- | --- | --- |
|  | **G** | **K** | **F** | **C** | **S** |
| **GKFCS** | 0.06 (0.02) | 0.12 (0.05) | 0.06 (0.03) | 0.38 (0.03) | 0.00 (0.02) |
| **GKFC** | 0.06 (0.02) | 0.12 (0.06) | 0.07 (0.03) | 0.38 (0.03) | * |
| **CAGE score** | | | | | |
|  | **G** | **K** | **F** | **C** | **S** |
| **GKFCS** | 0.15 (0.05) | 0.11 (0.15) | 0.00 (0.07) | 0.33 (0.08) | 0.00 (0.04) |
| **GKCS** | 0.16 (0.05) | 0.06 (0.08) | * | 0.30 (0.05) | 0.00 (0.04) |
| **GKC** | 0.17 (0.05) | 0.04 (0.07) | * | 0.31 (0.04) | * |
| **GC** | 0.19 (0.03) | * | * | 0.31 (0.04) | * |
| **Age onset smoking** | | | | | |
|  | **G** | **K** | **F** | **C** | **S** |
| **GKFCS** | 0.14 (0.05) | 0.06 (0.12) | 0.02 (0.06) | 0.07 (0.08) | 0.01 (0.03) |
| **GKCS** | 0.14 (0.05) | 0.10 (0.07) | * | 0.09 (0.05) | 0.02 (0.03) |
| **GKC** | 0.14 (0.05) | 0.12 (0.05) | * | 0.09 (0.05) | * |
| **Smoking Status** | | | | | |
|  | **G** | **K** | **F** | **C** | **S** |
| **GKFCS** | 0.20 (0.04) | 0.36 (0.09) | 0 (0.04) | 0.37 (0.05) | 0.10 (0.03) |
| **GKCS** | 0.22 (0.03) | 0.19 (0.05) | * | 0.29 (0.04) | 0.10 (0.03) |
| **Cigarettes per day** | | | | | |
|  | **G** | **K** | **F** | **C** | **S** |
| **GKFCS** | 0.20 (0.05) | 0.26 (0.12) | 0 (0.06) | 0.11 (0.08) | 0 (0.03) |
| **GKCS** | 0.20 (0.05) | 0.21 (0.07) | * | 0.08 (0.05) | 0 (0.03) |
| **GKC** | 0.21 (0.05) | 0.20 (0.06) | * | 0.09 (0.05) | * |

ST2) Showing full model selection using backward stepwise selection procedure in Generation Scotland.

|  | **Baseline** | | | **Age + Sex** | | | **Age + Sex + Test Centre** | | |
| --- | --- | --- | --- | --- | --- | --- | --- | --- | --- |
| **Trait** | **Beta** | **SE** | **P** | **Beta** | **SE** | **P** | **Beta** | **SE** | **P** |
| **Smoking Status**  **N=1653** | 0.19 | 0.02 | 6 x 10^-15^ | 0.21 | 0.02 | < 2 x 10^-16^ | 0.21 | 0.005 | < 2 x 10^-16^ |
| **Smoking Age Onset**  **N=449** | 0.05 | 0.04 | 0.24 | 0.04 | 0.04 | 0.35 | 0.04 | 0.04 | 0.36 |
| **Smoking Cigs per Day**  **N=381** | 0.08 | 0.05 | 0.12 | 0.11 | 0.05 | 0.04 | 0.10 | 0.05 | 0.05 |
| **Alc units per week**  **N=1406** | 0.26 | 0.03 | < 2 x 10^-16^ | 0.41 | 0.02 | < 2 x 10^-16^ | 0.40 | 0.02 | < 2 x 10^-16^ |
| **CAGE score**  **N=498** | 0.22 | 0.05 | 1 x 10^-6^ | 0.24 | 0.05 | < 2 x 10^-7^ | 0.23 | 0.05 | 4 x 10^-7^ |

**ST3**) Phenotypic associations between substance use phenotypes in opposite-sex couples in Generation Scotland (N=1742 total pairs). N shown in trait column reflect the N where both members of the couple had available phenotype data

|  | **All couples** | | | **Female couples** | | | **Male couples** | | |
| --- | --- | --- | --- | --- | --- | --- | --- | --- | --- |
| **Trait** | **Beta** | **SE** | **P** | **Beta** | **SE** | **P** | **Beta** | **SE** | **P** |
| **Smoking Status**  **N=278** | 0.24 | 0.06 | 5 x 10^-5^ | 0.16 | 0.08 | 0.05 | 0.31 | 0.08 | 3 x 10^-4^ |
| **Smoking Age Onset**  **N=53** | -0.11 | 0.12 | 0.35 | -0.16 | 0.21 | 0.46 | -0.10 | 0.16 | 0.53 |
| **Smoking Cigs per Day**  **N=49** | 0.50 | 0.16 | 0.003 | 0.73 | 0.24 | 0.005 | 0.31 | 0.22 | 0.17 |
| **Alc units per week**  **N=255** | 0.51 | 0.05 | < 2 x 10^-16^ | 0.50 | 0.07 | 2 x 10^-11^ | 0.52 | 0.09 | 1 x 10^-8^ |
| **AUDIT score**  **N=88** | 0.35 | 0.09 | 1 x 10^-4^ | 0.21 | 0.12 | 0.10 | 0.49 | 0.13 | 3 x 10^-4^ |
| **AUDIT-C**  **N=88** | 0.38 | 0.09 | 6 x 10^-5^ | 0.38 | 0.14 | 0.009 | 0.37 | 0.12 | 0.003 |
| **AUDIT-P**  **N=88** | 0.20 | 0.10 | 0.05 | 0.17 | 0.12 | 0.17 | 0.29 | 0.18 | 0.13 |

ST4) Phenotypic associations between substance use phenotypes in same-sex couples in UKB (407 pairs: 218 female pairs and 189 male pairs). N shown in trait column reflect the N where both members of the couple had available phenotype data and this is shown for the ‘All couples sample’.
